# Protein-Formatted Biomarker Enrichment Tag for the Detection of Carcinoembryonic Antigen

**DOI:** 10.1101/2025.01.15.633204

**Authors:** Chang Liu, Sara Barricella, Christina Cortez-Jugo, David L. Steer, Lizhong He

**Author notes:** Corresponding Authors Dr. Chang Liu, Department of Chemical and Biological Engineering, Monash University, Clayton, VIC 3800, Australia.,;, Dr. David Steer, Biomedicine Discovery Institute, Monash University, Clayton, VIC 3800, Australia.

## Abstract

The streptavidin-biotin interaction has long been a cornerstone of immunoassays, valued for its exceptionally high affinity and specificity, and has been widely used in routine diagnostic assays for decades. However, elevated biotin levels in plasma, often resulting from oral supplements, can interfere with streptavidin-biotin binding, compromising signal detection in these assays. To address this limitation, we present a biotin-insensitive antigen enrichment system based on the ultra-strong binding pair barnase (Bn) and barstar (Bs) for effective antigen capture. This high-affinity system enables efficient detection of nanobody-antigen complexes with picomolar sensitivity, as demonstrated by the detection of carcinoembryonic antigen (CEA, CEACAM5). When paired with a fluorescent secondary nanobody reporter, the system achieved a detection limit as low as 98.9 pM, with a limit of detection (LOD) of 17.8 ng/mL, representing a four-order-of-magnitude improvement in sensitivity compared to light interferometry-based binding analysis. Both Bn and Bs are small (<15 kDa), single-chain proteins that can be generically incorporated into immunoactive molecules. Compared to the traditional biotin-streptavidin system, the Bn-Bs system offers enhanced specificity, improved stability, and reduced cost and complexity.

## Introduction

Immunoassays (**IAs**) and diagnostic devices play a critical role in healthcare settings, enabling the detection of large and small biomolecules for disease diagnosis, drug monitoring, and other pharmaceutical analyses. IAs rely on the specific binding of an antibody (**Ab**) to its cognate antigen (**Ag**). Most antibodies used in IAs exhibit binding affinities, measured by the equilibrium dissociation constant (**K**_**D**_), in the nanomolar range (10^-7^ to 10^-9^ M) range^1^. However, *in vitro* diagnostic practices often require the detection of biomarkers (antigens) at concentrations as low as pg/mL to ng/mL, corresponding to femtomolar (fM) to picomolar (pM) levels for typical protein biomarkers with molecular weights ranging from 50 to 200 kDa^2^. To achieve pM (or sub-nanomolar) detection of biomarkers using nanomolar-affinity antibodies, a two-step amplification strategy is commonly employed. This approach involves first enriching the Ab-Ag complex to a higher local concentration, followed by amplifying the signal derived from reporter molecules.

A well-established technique for antigen enrichment utilizes streptavidin (SA)-coated nanoparticles (NPs) and a “retrievable” biotin-labelled primary monoclonal antibody. The biotin-labelled primary antibody is pre-mixed with the antigen, and upon the addition of streptavidin-coated NPs, the strong biotin-streptavidin interaction (K_D_ = 10^−15^ M) ^3^ facilitates the enrichment of the antigen on the NP surface. This ensures stable adsorption of the antibody-antigen complex to the solid surface with high loading capacity, effectively concentrating picomolar-level biomarkers into a detectable range. Subsequently, an antibody-antigen-antibody complex is formed by binding a reporter secondary antibody, enhancing detection sensitivity and reproducibility. The biotin-streptavidin interaction is the foundation of several commercial diagnostic systems, such as the Roche Elecsys® immunochemistry platform^4^.

Although the biotin-streptavidin system is employed in over 50% of immunoassays due to its outstanding binding kinetics and reliability, it can be susceptible to interference from biotin components present in the analyte^5^. Biotin (also known as Vitamin B7, Vitamin H, or Coenzyme R) naturally occurs in human serum and urine^6^, where it plays a critical role in metabolism as a cofactor for carboxylase enzymes. The growing trend of overconsumption of biotin supplements has led to interference in a wide range of immunoassays, including those for thyroid markers^7^, hormones^8^, and the cardiac function biomarkers^9^. Excess amount of biotin competes with the formation of sandwich immune complexes and resulting in false positives. Current interference mitigating methods, including monitoring serum biotin level^10^, controlled biotin intake^8^, and removal of biotin prior to the assay^11^, inflict complications and uncertainty on the standard assay Another limitation of the biotin-streptavidin system is its high production cost, primarily due to the requirement for *in vivo* or *in vitro* biotinylation of the primary antibody^9^, and the relatively low productivity of streptavidin expression (100-120 mg/L in 8-10 day^12^).

Herein, we present the design of an alternative antigen enrichment method that can be generically applied to biosensing applications. This approach leverages the high specificity and strong affinity of the protein-based binders barnase (Bn) and barstar (Bs), originally identified from *Bacillus amyloliquefaciens*^12^. In nature, barnase functions as an extracellular ribonuclease that is cytotoxic to *Bacillus amyloliquefaciens*, thus, it must be co-expressed with its inhibitor, barstar, prior to secretion^13^. The selection of Bn and Bs is guided by key factors critical to immunochemistry. First, Bn and Bs exhibit exceptionally tight binding (K_D_ of 10^-14^ M)^12^, comparable to the benchmark streptavidin and biotin interaction (K_D_ of 10^-15^ M), allowing effective capture of the antigen, even at low concentration, into the sandwich complex. Second, Bn and Bs are proteins forms stable one-to-one (Bn-Bs) complex^13^, inherently eliminating the risk of biotin interference. Finally, barnase and barstar are small proteins (12 kDa and 10 kDa, respectively) with robust structural stability, making them ideal candidates for use as protein tags.

## Results and discussion

Most SA-biotin systems adapt the strategy to coat the surface with streptavidin to capture biotinylated antibodies (**Scheme 1a**). This approach is primarily due to the large-size and multimeric nature of streptavidin (66 kDa, tetramer), which limits its use as a tagging protein. Attempts in engineering monomeric streptavidin often results in lower binding affinity^14^ or increased production complexity^15^. Popular antigen enrichment technologies use solid support such as nanoparticles (NPs) for increased ligand density and ease of isolation^4^.

The primary consideration in designing a Bn-Bs-based detection system is ensuring compatibility with the charge characteristics of the fusion protein candidates. To demonstrate the functionality of the devised system, we recombinantly fused Bn/Bs with a pool of nanobodies. The tight interactions between Bn and Bs are well-understood, with one key determining factor being the electrostatic interaction, as Bn (isoelectric point, pI=8.8) and Bs (pI=4.6) exhibit opposite charges under physiological pH conditions^16^. This poses a challenge in designing Bn/Bs and Nb fusions, as oppositely charged proteins tend to be less stable. To address this, two complementary strategies for protein capture are proposed: (1) the use of a barnase-coated matrix to capture Bs-fused, negatively charged nanobodies, and (2) the use of a barstar-coated matrix to capture Bn-fused, positively charged nanobodies (**Scheme 1b, c**). These approaches aim to improve the stability of the chimeric protein constructs.

A significant challenge associated with the Bn-Bs system is the cytotoxic, ribonuclease nature of barnase. Expression of Bn or Bn-tagged recombinant protein is lethal to the host, unless the barnase is removed from the cytoplasm with its inhibitor barstar co-expressed^17^. We first expressed barnase using a plasmid pMT416 which includes a phoA signal peptide for the re-direction of the expressed peptides to *E. coli* periplasm^18^. The barnase protein was successfully purified in accordance with previously established methods^18^ using lysis-free sucrose extraction (**Supplementary Figure S2 a**).

Two nanobody candidates, 2D5 (primary) and 13A5 (secondary), which can bind CEA simultaneously, were selected^19^. We constructed two chimeric proteins—2D5-Bn and 2D5-Bs. Given that the primary nanobody 2D5 has a pI of 8.66, the Bn tag is more likely to remain stable at physiological pH. A (G4S)2 flexible linker, with an estimated length of 4 nm, was incorporated to enhance proper folding. The secondary nanobody, 13A5, was conjugated with NHS-ATTO488 for straightforward fluorescent signal reporting. Although fluorescent intensity can be influenced by instrumental settings, such as laser intensity and photomultiplier power, these parameters can typically be standardized, enabling the comparison of relative intensities under controlled conditions.

To express the chimeric 2D5-Bn fusion, two different approaches were explored: (1) dual transformation of 2D5-Bn in pBAD33 (chloramphenicol resistant, arabinose-inducible pBAD promoter) and Bs in pET28a (kanamycin resistant, T7 promoter) in the host cell *E. coli* BL21(DE3) (**Supplementary Figure S1**); and (2) co-expression of 2D5-Bn and its inhibitor Bs in a single plasmid (pNbBnBs) using the pETDuet-1 backbone (**Figure 1d**). Under double selection pressure (chloramphenicol and kanamycin), the dual-transformation strategy exhibited low transformation efficiency and was outperformed by the successful transformation of the pETDuet-1 plasmid encoding both 2D5-Bn and Bs.

**Figure 1.**
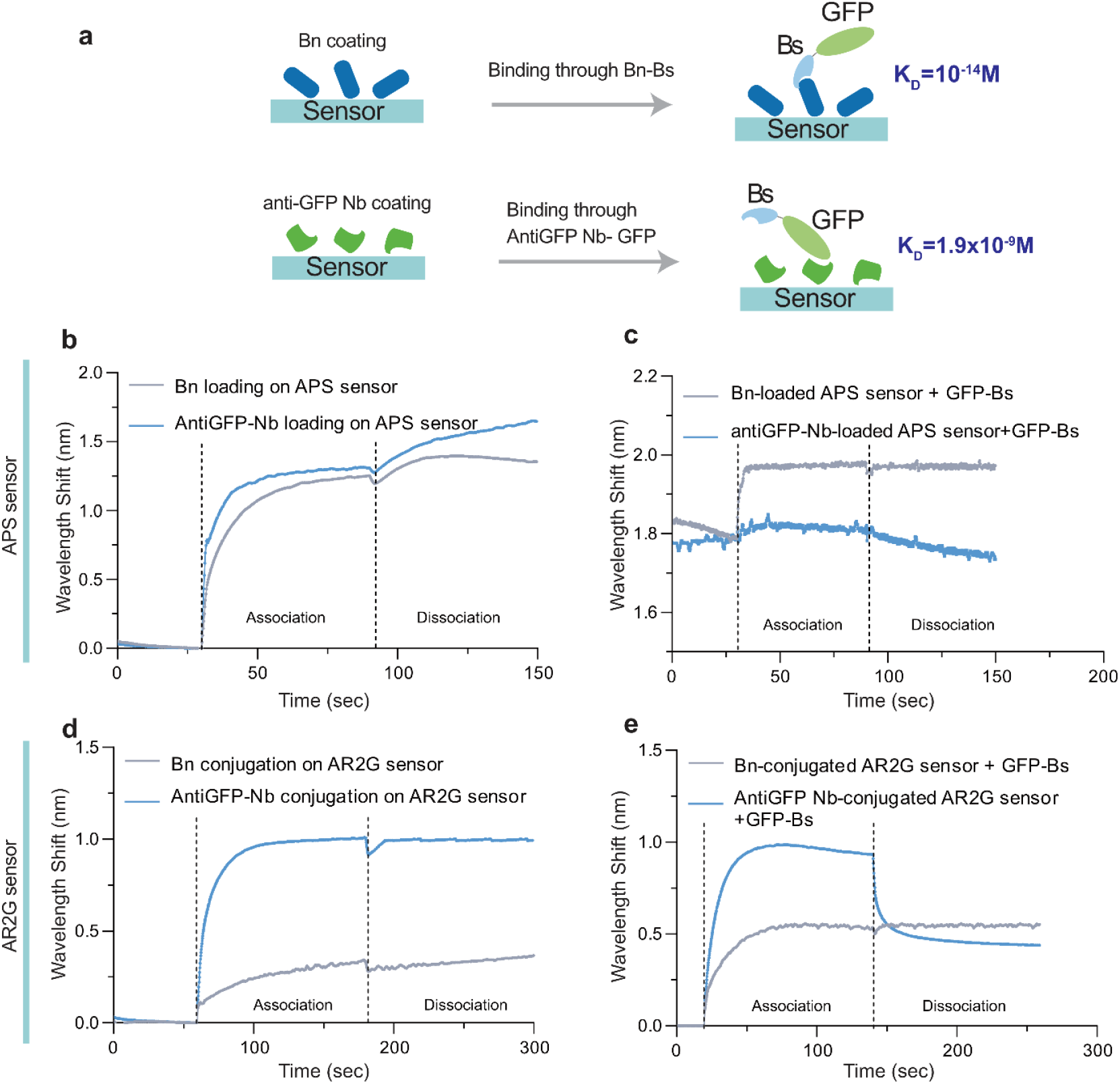
Comparison of Bn-Bs interaction and anti-GFP nanobody-GFP interaction on APS- and AR2G-coated sensors using light interferometry. **a**, Illustration of surface interactions facilitated by Bn-Bs and anti-GFP-Nb and GFP. **b**, Passive adsorption of Bn and anti-GFP Nb on an APS sensor (representative curves from n=2). **c**, GFP-Bs capture on Bn-coated and anti-GFP-coated APS sensors (representative curves from n=2). The K_D_ for Bn–Bs-GFP is below the quantifiable range, while the K_D_ for anti-GFP Nb–GFP-Bs cannot be determined due to unstable adsorption. **d**, Chemical conjugation of Bn and anti-GFP Nb on an activated AR2G sensor (representative curves from n=2). **e**, GFP-Bs capture on Bn-conjugated and anti-GFP-conjugated AR2G sensors (representative curves from n=2).

The pBnBS1 plasmid enables simple periplasmic expression of 2D5-Bn in rich media (**Supplementary Figure S2c**) using the host *E. coli* SHuffle, which enhances cytoplasmic formation of disulfide bonds^20^. After purification, all proteins remained stable at 4 °C, except for 2D5-Bs, which exhibited precipitation after one week of incubation at 4 °C. Desalting 2D5-Bs into 20 mM Tris-HCl (pH 8) with 140 mM NaCl immediately after affinity chromatography purification effectively stabilized the protein. The identity of the purified proteins was confirmed by mass spectrometry. Peptide matching demonstrated high-confidence sequence coverage (**Supplementary Table S3**) and confirmed the auto-truncation of the phoA periplasmic signal peptide for periplasmic phoA-Bn (bare Bn consists of 110 amino acid residues).

**Scheme 1.**
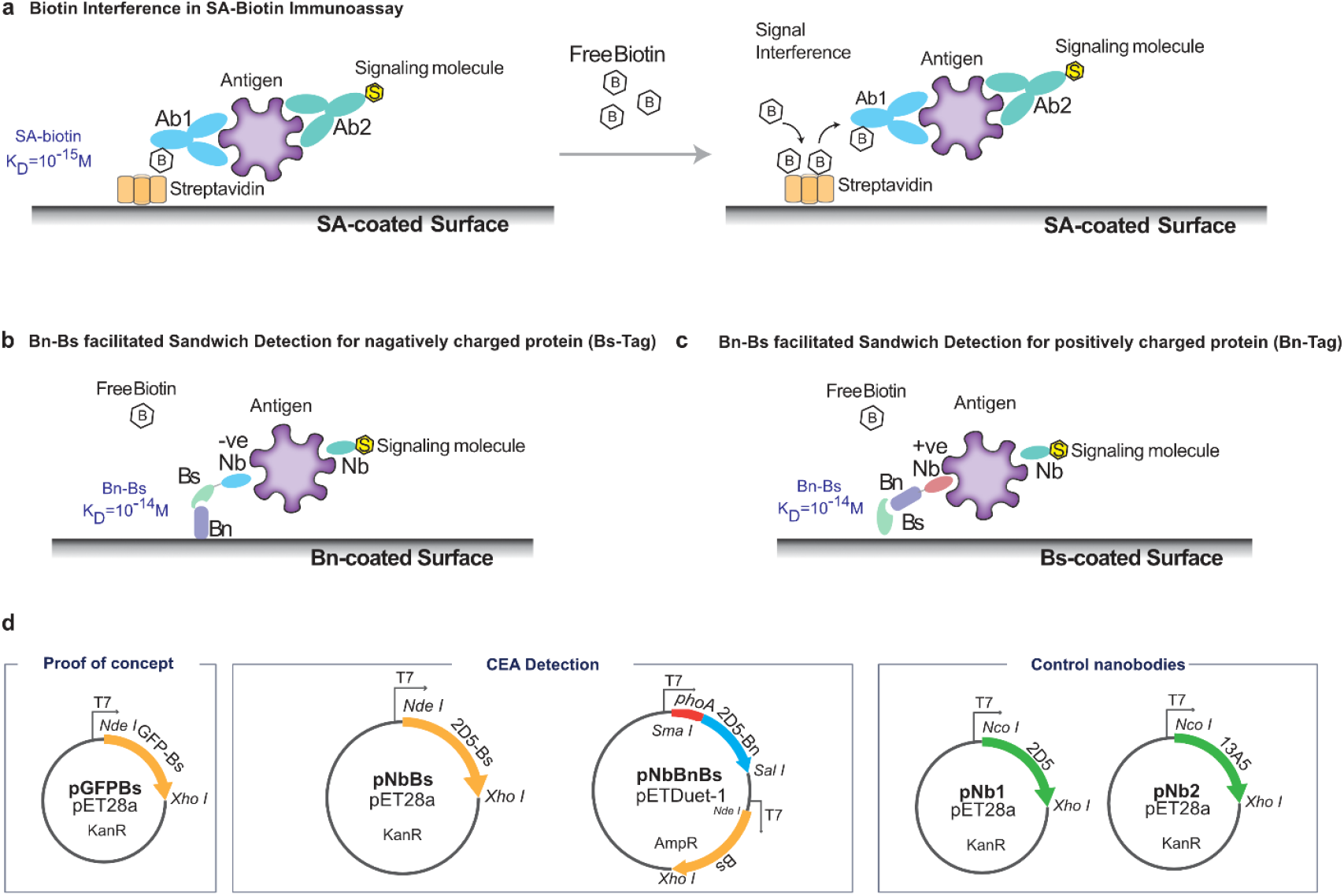
Design of protein-formatted biomarker enrichment strategy using barnase (Bn) and barstar (Bs) Tags. **a**, The SA-biotin system, which uses streptavidin-labelled beads to capture the Ab-Ag complex, is sensitive to biotin interference. **b**, Antigen enrichment with Bs-tag for proteins (e.g. Nb, nanobodies) that are negatively charged at physiological pH (pI<7.4). Bs-tagged protein can be enriched on Bn-coated surface. **c**, Antigen enrichment with Bn-tag for proteins (e.g. Nb, nanobodies) that are positively charged at physiological pH (pI>7.4). Bn-tagged protein can be enriched on Bs-coated surface. **d**, Plasmids constructed in this study. From left to right: Plasmid encoding GFP-Bs; Plasmids for CEA detection encoding 2D5-Bs and 2D5-Bn; Plasmids encoding nanobodies 2D5 and 13A5 as control.

In an effort to understand the binding behaviour of barnase (Bn) and barstar (Bs) at solid interfaces, a green fluorescent protein-Bs (GFP-Bs) fusion protein was synthesized as a model molecule (**Scheme 1d**), along with an anti-GFP nanobody (Nb) LaG-14^21^ (Addgene #172745, K_D_ =1.9×10^-9^ M) to allow dynamic tracking of the Bn-Bs interactions using a light interferometer (BLItz, Pall ForteBio). The binding dynamic between Bn-Bs and the nanobody-GFP was investigated first on aminopropyl silane (**APS**) surface sensors. Two groups of APS sensors were saturated with excess Bn or anti-GFP Nb through non-covalent, hydrophobic adsorption (**Figure 1 b**), respectively. After washing away unbound proteins with phosphate-buffered saline (PBS), 2.5 µM of GFP-Bs in 3% bovine serum albumin (BSA) solution was introduced. As expected, the APS sensor coated with Bn exhibited high affinity toward GFP-Bs, with a K_D_<10^-10^ M, which is below the quantifiable range of the BLItz instrument. In contrast, the anti-GFP Nb, despite having a reported KD in the nanomolar range, failed to effectively capture the GFP molecule (**Figure 1b, c**).

To elucidate the effect of ligand density at interface on the binding behavior, the experiment was repeated on carboxyl surface (Octet® AR2G, Sartorius) sensors for controlled covalent conjugation. Following the activation of surface carboxyl group using carbodiimide linker (EDC/sNHS), two groups of sensors are loaded with equimolar amount (4 uL of 25 uM) of Bn and anti-GFP Nb, respectively. Both Bn and anti-GFP Nb were successfully conjugated to the surface carboxyl group, as evidenced by the negligible dissociation phase (**Figure 1d**). Notably, Bn exhibited a lower biolayer thickness compared to the anti-GFP Nb (**Figure 1d**), which is expected given that Bn contains fewer reactive amine groups than GFP-Bs (8 lysine residues versus 26 lysines in GFP-Bs). After quenching the excess NHS with ethanolamine, 2.5 µM of GFP-Bs in PBS was subsequently introduced to the sensors. The capture of GFP-

Bs by Bn yielded a dissociation constant that fell below the instrument’s quantifiable range, further confirming the high affinity of the Bn-Bs interaction observed on the APS sensor. Conversely, the immobilized anti-GFP Nb rapidly captures GFP-Bs from the solution but exhibits a fast wash-off of the captured GFP-Bs during the dissociation phase (**Figure 1e**). Collectively, these results demonstrate that the strong affinity of the Bn-Bs binding pair enables the effective capture of biomolecules at the solid interface.

We next investigated protein capture efficiency on microbeads, selecting a well-defined surface—NHS-activated Sepharose microspheres (dV50=90 μm)—for covalent bioconjugation (**Figure 2a**). Each sample was prepared using a known volume of NHS beads capable of capturing approximately 0.66 nmol of protein^22^. The bead surface was saturated with a high concentration of either Bn or anti-GFP Nb, and unreacted NHS residues were quenched with 1 M Tris-HCl (pH 8). Surprisingly, rapid saturation of the anti-GFP ligand was observed: assuming a one-to-one binding ratio for both the Bn-Bs and anti-GFP nanobody-GFP systems, an estimated 12.8 µg of GFP-Bs would be required to saturate 50% (0.33 nmol) of the surface-bound Bn. However, fluorescence measurements revealed that the anti-GFP Nb reached saturation after the addition of 50 µL of 250 nM GFP-Bs (corresponding to 475 ng; **Figure 2b**). In contrast, Bn-coated microspheres exhibited a distinct semi-linear response to increasing GFP-Bs concentrations, underscoring the efficacy of the Bn-Bs system in enhancing surface protein capture from biofluids. Additionally, the Bn-Bs binding interaction remained stable and was unaffected by the presence of biotin, even at concentrations as high as 3.2 µM (**Figure 2c**).

**Figure 2.**
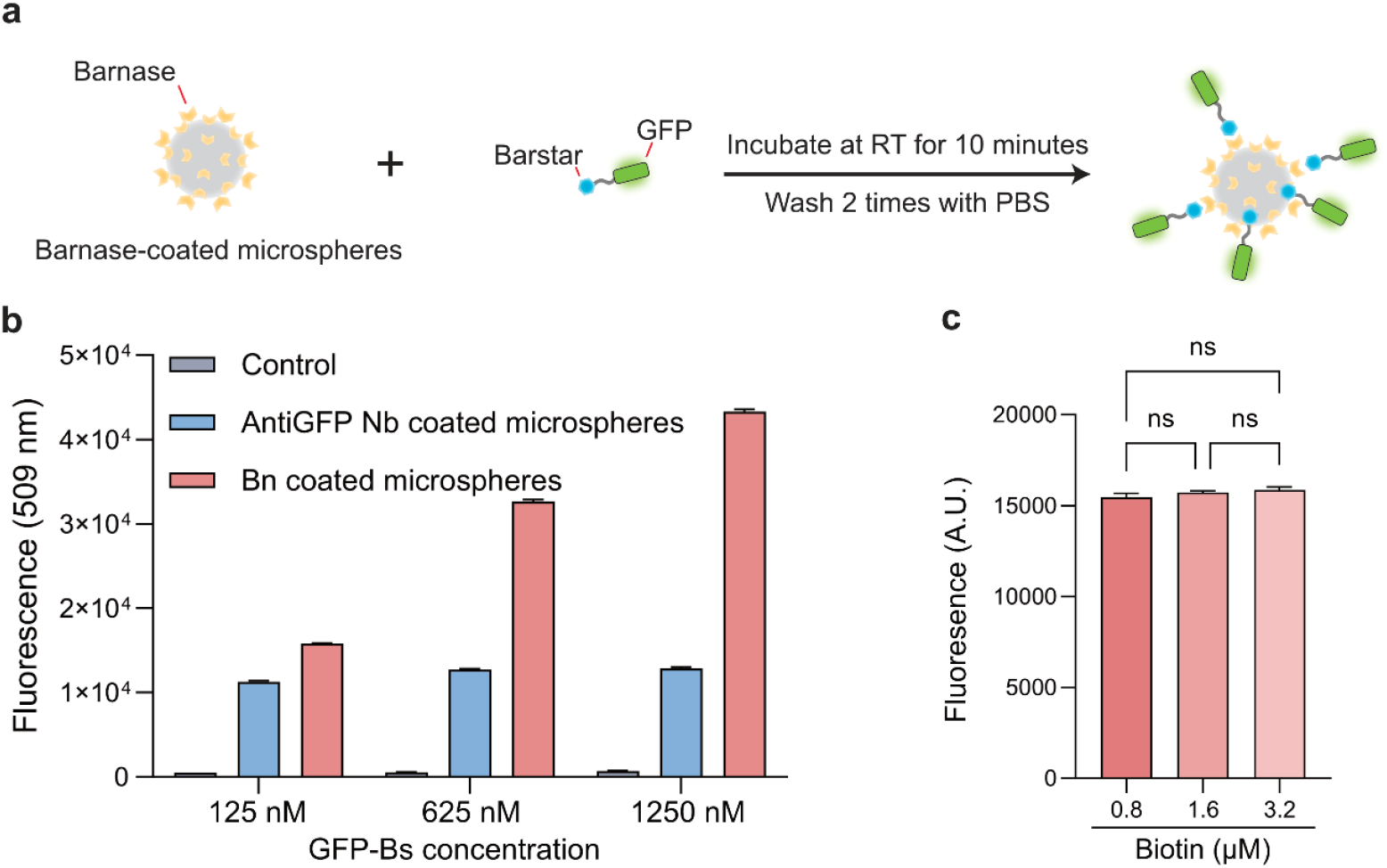
Bead-based antigen capture using the Bn-Bs enrichment system. **a**, Bn-conjugated Sepharose microspheres were used to capture GFP-Bs in PBS containing 3% BSA. After incubation and washing, GFP fluorescence of the beads was measured at an emission wavelength of 520 nm. **b**, Analysis of protein capture capacity of Bn and anti-GFP Nb-conjugated beads. Fluorescence was measured for washed microspheres incubated with 50 µL of GFP-Bs at final concentrations of 125 nM (475 ng), 625 nM (2.38 µg), and 1250 nM (4.75 µg), respectively (total reaction volume: 100 µL). The maximum loading capacity of the conjugated beads was estimated at 12.8 µg of GFP-Bs. **c**, Biotin has a negligible effect on GFP-Bs capture by Bn-conjugated microspheres. Biotin was added to the capture reaction at final concentrations of 0.8 µM, 1.6 µM, and 3.2 µM, where 20 µL of beads was used to capture 475 ng (125 nM final concentration) of GFP-Bs.

The effective surface capture enabled by the Bn-Bs interaction facilitated the further development of a sensing system designed for the detection of human carcinoembryonic antigen (**CEA**). Two complementarily nanobodies (Nbs, also denoted as V_H_H) against CEA were selected^19^. Nanobodies have shown distinct advantages such as superior stability and comparable binding capacities to conventional antibodies or antibody fragments while possessing significantly smaller molecular weight (∿15 kDa)^23^. Despite the identification of nanobodies against a range of antigens^21,24,25^, the commercial availability of nanobody-based biosensors remains limited due to several technical challenges. First, immobilized nanobodies often adopt random orientations, which can hinder access to their binding sites. This contrasts with full-length antibodies, which are typically conjugated via the Fc domain, ensuring a more uniform orientation. Second, the chemical or enzymatic conjugation of biotin or other functional molecules to nanobodies—significantly smaller in size compared to full-length antibodies—frequently results in a reduction in binding affinity, a challenge less pronounced with full-length antibodies. Fortunately, these limitations can be addressed by recombinantly fusing nanobodies with the Bn-Bs system. Owing to their small size, nanobodies can be seamlessly fused to the Bn/Bs tagging system, producing fusion proteins of approximately 30 kDa that can be efficiently expressed in a single step using cost-effective prokaryotic hosts. Additionally, the incorporated Bn-Bs system enables one-to-one, site-specific capture of nanobodies, maximizing the exposure of their binding sites for effective antigen interaction.

The performance of 2D5 and 13A5 was subsequently benchmarked for the enrichment of CEA using a previously developed silica-binding peptide, CotB1p, incorporated into 2D5 on a regenerated quartz sensor surface, following a previously established method^26^. Both 2D5 and 13A5 demonstrated affinity toward CEA, as shown in **Supplementary Figure S5**. However, this configuration was limited to detecting CEA at concentrations above 0.1 µM (18 µg/mL).

Identification of CEA within the normal plasma concentration range (0-2.5 ng/mL in non-smokers and 0-5 ng/mL in smokers^27^) holds significant diagnostic and prognostic value for colorectal^27^, breast^27^, lung^28^ and pancreatic^29^ cancer. Therefore, the capability to detect CEA at picomolar (low nanogram/mL) levels is of substantial clinical relevance.

To improve the detection of CEA, a simple Nb-Ag complex capture assay using the Bn-Bs system was designed (**Figure 3a**). Briefly, the assay cocktail consisted of the primary and secondary nanobody, i.e., 2D5-Bn and 13A5-ATTO488, respectively. After thoroughly mixing the assay cocktail with CEA, the bound Nb-CEA-Nb complex was enriched by adding Bn-coated microspheres. In the final step, the microspheres were isolated, washed, and their fluorescence was measured using fluorescence spectroscopy.

**Figure 3.**
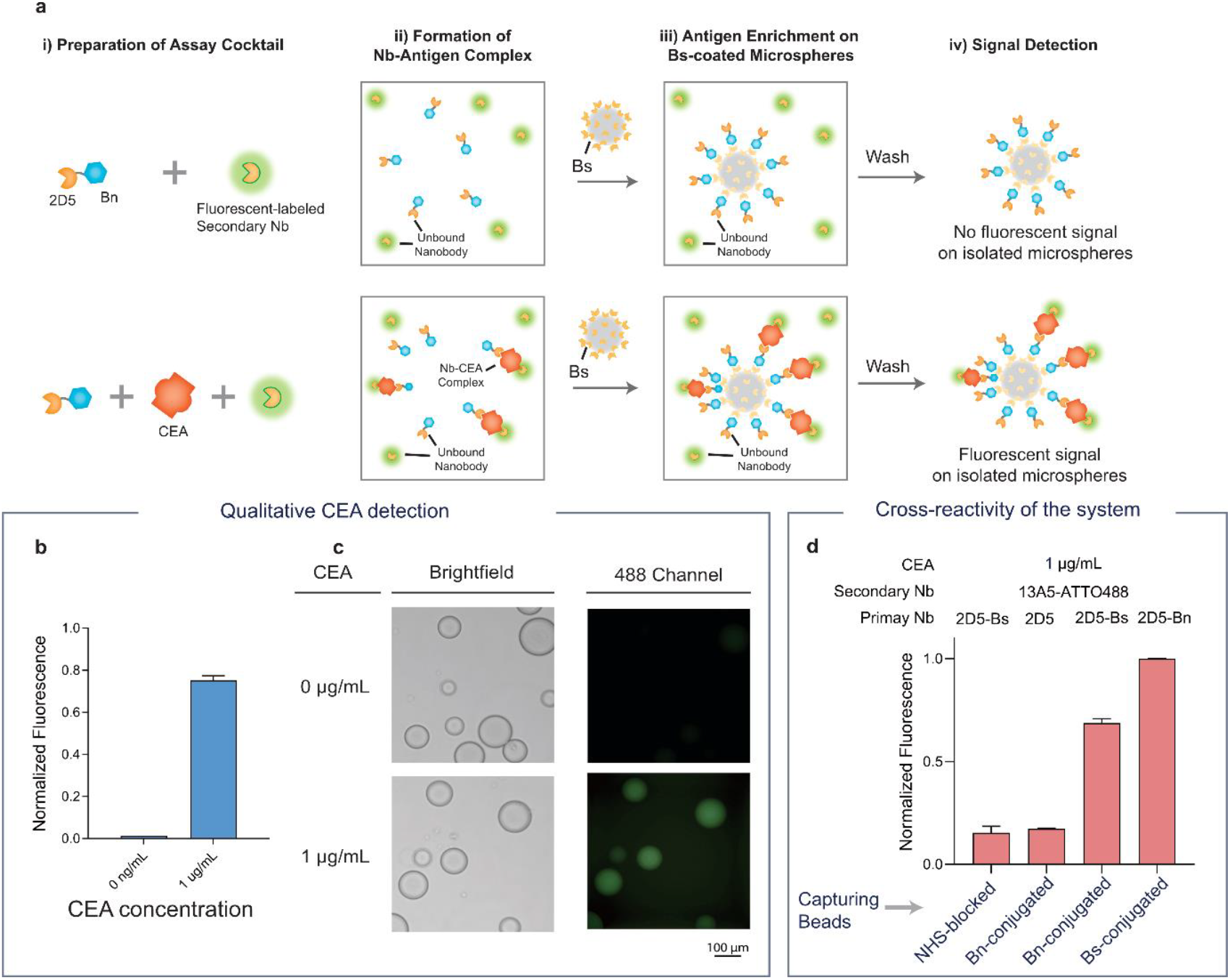
Detection of CEA using anti-CEA nanobodies 2D5 and 13A5. **a**, Illustration of CEA detection with the Bn-Bs Nb-antigen enrichment system. Bs-conjugated NHS-activated Sepharose microspheres were used for the isolation of the 2D5-Bn–CEA complex. The isolated microspheres were washed twice with PBS prior to fluorescence measurement. **b**, Qualitative CEA detection using the proposed method. The standard detection reaction was set up with 20 µL of 0.1 mg/mL 2D5-Bn, 20 µL of CEA at 1 µg/mL, and 20 µL of 0.1 mg/mL 13A5-ATTO488. After incubation in the dark for

To minimize variations in fluorescence signals due to exposure and sensor settings, data acquisition was standardized and kept consistent using a Tecan Infinite Pro-200. An initial assessment was conducted to evaluate the signal viability and feasibility for detecting low nanomolar concentrations (1 µg/mL) of CEA using oversaturated 2D5-Bn and Bs-coated beads. Under standard detection conditions (20 µL of 0.1 mg/mL 2D5-Bn, 20 µL of CEA at 1 µg/mL, and 20 µL of 0.1 mg/mL 13A5-ATTO488 captured with 10 µL of 50 w/w% Bs-conjugated beads), the system yielded a distinct signal for 1 µg/mL CEA, while the control group (assay cocktail without CEA) exhibited baseline intensity (**Figure 3b**). This observation was corroborated by visualizing the isolated microspheres using a fluorescence microscope (**Figure 3c**).

Subsequently, the specificity of the designed system was assessed. The fluorescence signals generated from (1) 2D5-Bn and Bs-facilitated enrichment, (2) 2D5-Bs and Bn-facilitated enrichment, (3) non-specific 2D5 and Bn interaction, and (4) non-specific Tris-blocked microspheres and 2D5-Bs were compared using an excess amount of primary and secondary nanobodies (20 µL of 0.5 mg/mL primary Nb and 20 µL of 0.5 mg/mL 13A5-ATTO488) in phosphate-buffered saline with Tween 20 (PBST) containing 3% BSA per test. Although a distinct signal was observed for 2D5-Bn and Bs, elevated fluorescence (approximately 15% higher) was detected for non-specific surface interactions (**Figure 3d**), which was not observed in the CEA-absent control (**Figure 3b**). This relatively high noise level is unlikely due to non-specific Bn/Bs interactions with the test protein, as the Bn-coated surface and Tris-blocked surface exhibited similar fluorescence levels. Instead, the non-specific signal is likely caused by weak interactions between the multimeric, glycoprotein-formatted CEA and agarose-based microspheres. Notably, 2D5-Bs captured with Bn-coated particles showed a weaker signal compared to 2D5-Bn captured with Bs-coated particles, potentially due to the reduced stability of the chimeric 2D5-Bs fusion (**Figure 3d**).

10 minutes, the Nb-CEA complex was captured with Bs-conjugated beads. **c**, Fluorescence microscopy of isolated microspheres incubated in the absence or presence of CEA. Images were taken using the same settings. **d**, Evaluation of non-specific interactions between detection components. The detection reaction was set up with primary and secondary nanobodies in excess—20 µL of 0.5 mg/mL primary Nb, 20 µL of CEA at 1 µg/mL, and 20 µL of 0.5 mg/mL 13A5-ATTO488. NHS-blocked beads had their NHS groups deactivated with 1 M Tris-HCl (pH 7). The fluorescence baseline for **b** and **d** was set to the PBS signal.

Ultimately, we aimed to establish a quantitative correlation between the fluorescence signal and the concentration of CEA. The fluorescence signal exhibited a Langmuir adsorption correlation with CEA concentration (**Figure 4a, b**). At low CEA concentrations (10–100 ng/mL), a linear correlation was observed between fluorescence intensity and CEA concentration. The limit of detection (LOD) was calculated to be 17.8 ng/mL using Equation 1, corresponding to 98.9 pM of CEA. Compared to the micromolar-level detection achieved with planar surface-immobilized 2D5 (**Supplementary Figure S5**), the Bn-Bs antigen enrichment system enabled a four-order-of-magnitude improvement in detection sensitivity, using a simple fluorescent reporter molecule.

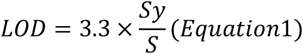

where Sy is the standard error of linear regression and S is the slope

These findings demonstrate that the Bn-Bs antigen enrichment system, combined with a simple fluorescent reporter, represents a transformative approach for ultrasensitive biomarker detection, offering a robust and cost-effective platform with significant potential for clinical diagnostics and early disease detection.

**Figure 4.**
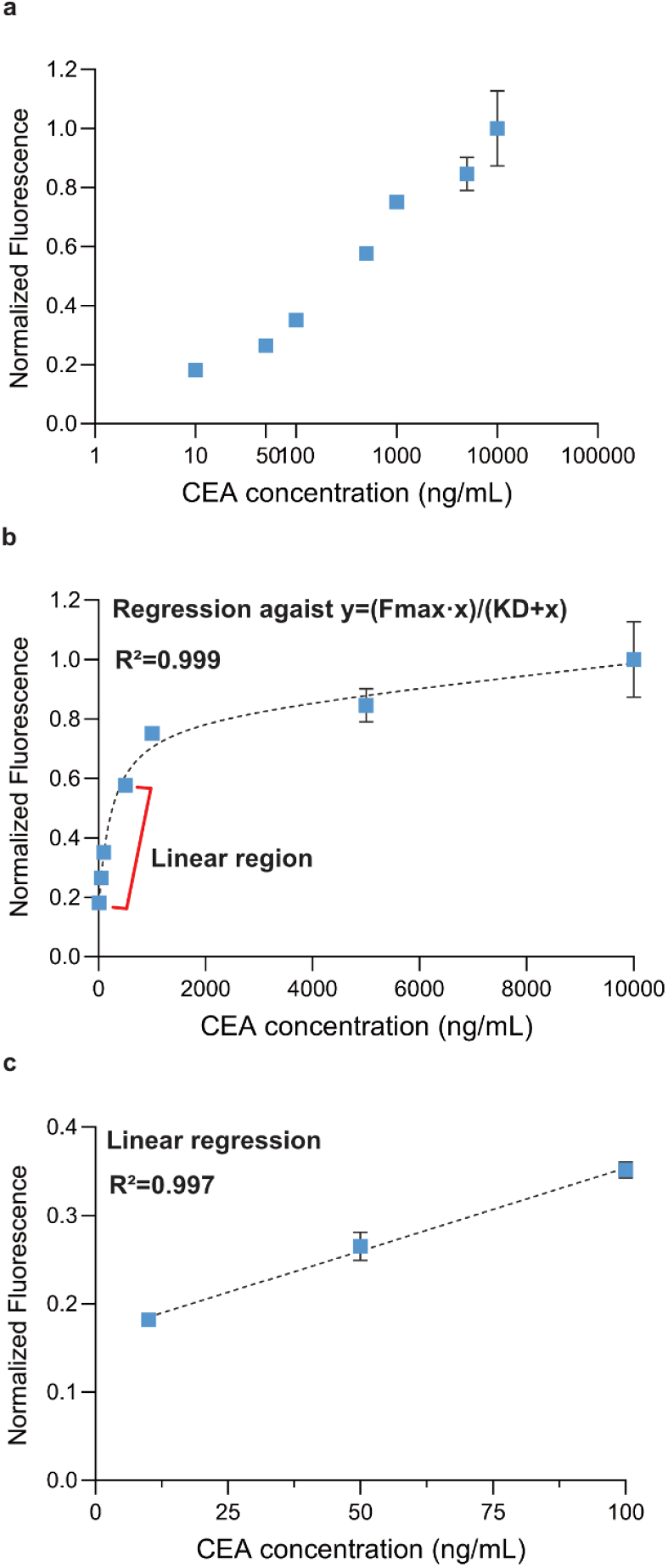
Quantitative analysis revealing the potential for low picomolar detection of CEA. **a**, Fluorescence signal of detected CEA under standard condition: 20 μL of 0.1 mg/mL primary Nb, 20 μL of 0.1 mg/mL 13A5-ATTO488, 10 µL of 50 w/w % conjugated beads for the detection of 20 μL CEA at various concentration (from 10-100,000 ng/mL). **b**, Non-linear regression indicated a Langmuir adsorption correlation (one site specific binding). Apparent K_D_ was estimated at 244.1 ng/mL for Nb-CEA complex, equivalent to approximately 976 pM (Nb-CEA-Nb of around 250 kDa). **c**, Linear correlation at low CEA concentration revealed a limit of detection (LOD) of 17.8 ng/mL.

## Conclusions

In this work, we report an alternative protein-formatted approach for antigen enrichment in detection assays using the ultra-strong binding pair Bn and Bs. Given that Bn and Bs exhibit specific one-to-one binding, the system provides a fundamental solution to the challenge of biotin interference. Through systematically designed studies, the Bn-Bs-enabled antigen enrichment system demonstrated a four-order-of-magnitude improvement in the detection of CEA, achieving a limit of detection (LOD) of 17.8 ng/mL (98.9 pM). We anticipate that even higher sensitivity can be achieved by further optimizing the reporting system, such as through the use of sophisticated signal amplification methods like enzymatic or chemiluminescence reporting. Overall, the Bn-Bs enrichment system can be generically applied to a wide range of protein detection applications and enables rapid, low-cost production using prokaryotic hosts.

