## Supplementary Information for "Protein-Formatted Biomarker Enrichment Tag for the Detection of Carcinoembryonic Antigen"

* Corresponding Authors

Dr. Chang Liu

Department of Chemical and Biological Engineering, Monash University, Clayton, VIC 3800, Australia.

Dr. David Steer

Biomedicine Discovery Institute, Monash University, Clayton, VIC 3800, Australia

#### Material and method

*Material*

The plasmids used in this study are summarized in **Supplementary Table S1**. The plasmids encoding barnase (pMT416) and barstar (pMT316) were kindly provided by Robert Hartley (Addgene plasmids #8607 and #8608). The anti-GFP nanobody LaG-16 was a gift from Michael Rout (Addgene plasmid #172745). The amino acid sequences of the anti-CEA nanobodies 2D5 and 13A5 were generously provided by Prof. Rui Nian. Expression plasmids encoding Bn, Bs, GFP-Bs, 2D5-Bs, and 2D5-Bn were synthesized by GenScript (Hong Kong), and their sequences are detailed in **Supplementary Table S2**. Human carcinoembryonic antigen (CEACAM5) was purchased from Abcam (ab742). NHS-activated Sepharose 4 Fast Flow microspheres (product #17090601) were obtained from GE Healthcare. ABTS and NHS-ATTO488 dye were purchased from Sigma-Aldrich.

*Production of Bn containing constructs*

The expression of Bn coded pMT416 according to previous method1. For the expression of 2D5-Bn, two methods were attempted: 1) The pBAD33 plasmid encoding 2D5-Bn was transformed into *E. coli* BL21(DE3) by heat shock at 42 °C for 5 minutes. The transformants were incubated in SOC media for 1 hour and then plated on TB plates containing 50 μg/mL kanamycin and 25 μg/mL chloramphenicol. 2) The pETDuet-1 plasmid encoding 2D5-Bn and Bs was transformed into *E. coli* BL21(DE3) and DH5α. Overall, the double transformation resulted in low efficiency, with only 1–2 colonies observed for BL21(DE3). In contrast, the transformation of pETDuet-1 was successful and was therefore selected as the preferred method for 2D5-Bn expression.

Bn and 2D5-Bn were auto-induced in BL21(DE3) without additional inducers or heat induction. Protein expression was carried out in TB media at 37 °C for 12 hours prior to cell harvest. Both Bn and 2D5-Bn were periplasmically expressed, and ice-cold sucrose was added to a final concentration of 15% (w/v) to enhance protein recovery. Host cells were removed by centrifugation at 10,000 rpm and 4 °C for 15 minutes, and the supernatant was collected for further purification.

For Bn purification, the media containing pMT416-expressed Bn was diluted threefold with Milli-Q water before loading onto a 1 mL HiTrap SP HP cation exchange column (GE Healthcare). Bn was collected using gradient elution with Buffer A (20 mM Tris-HCl, pH 8) and Buffer B (20 mM Tris-HCl, pH 8, 1 M NaCl).

2D5-Bn was purified using a 1 mL HisTrap affinity column (GE Healthcare). Bound protein was eluted with an imidazole gradient using Buffer A (20 mM Tris-HCl, pH 8, 500 mM NaCl) and Buffer B (20 mM Tris-HCl, pH 8, 500 mM NaCl, 500 mM imidazole). Purified Bn and 2D5-Bn were desalted using a desalting column (GE Healthcare) into DPBS for long-term storage.

*Expression and purification of other proteins*

All proteins lacking the Bn moiety were expressed in rich Terrific Broth (TB). Plasmids encoding the nanobodies LaG-16, 2D5, 2D5-Bs, and 13A5 were transformed into *E. coli* SHuffle T7. The expression cultures, inoculated at an OD of 0.1, were incubated at 30 °C for 3 hours, followed by induction with 1 mM IPTG for 20 hours at 20 °C. For Bs and GFP-Bs, *E. coli* BL21(DE3) was used, and expression was induced with 1 mM IPTG, followed by culturing for 20 hours at 30 °C.

To harvest the proteins, cell cultures were centrifuged at 10,000 rpm and 4 °C for 10 minutes. The isolated cells were lysed, and cell debris was removed by centrifugation at 10,000 rpm and 4 °C for 15 minutes. Cell lysis was performed by sonication using lysis buffer (20 mM Tris-HCl, pH 8, 0.5 M NaCl, 0.5% Triton-X) for LaG-16, 2D5, 2D5-Bs, 13A5, GFP-Bs, and Bs.

Purification of LaG-16, 2D5, 2D5-Bs, 13A5, GFP-Bs, and Bs was carried out using an AKTA FPLC system (GE Healthcare), following a previously established method[2]. Briefly, the proteins were loaded onto HisTrap columns (GE Healthcare) and eluted with a 0–500 mM imidazole gradient in 20 mM Tris-HCl (pH 8) and 0.5 M NaCl. Purified proteins were desalted into DPBS for storage. For improved stability, 2D5-Bs was desalted into 20 mM Tris-HCl (pH 8) and 140 mM NaCl.

*Evaluating binding performance using Bn and antiGFP-Nb coated surface*

Two types of light interferometer sensors were used: APS sensors and amine-reactive AR2G sensors.

For APS sensors, after hydration in PBS at room temperature for 10 minutes, the sensors were equilibrated with 4 µL of 25 µM Bn or anti-GFP Nb, respectively. The coated sensors were washed with PBS, and 2.5 µM GFP-Bs in PBS containing 3% BSA was introduced to determine binding kinetics. Dissociation was measured by placing the sensor in PBS with 3% BSA.

For amine-reactive AR2G sensors, after hydration in PBS at room temperature for 10 minutes, the sensors were activated with freshly prepared 20 mM EDC and 10 mM sulfo-NHS at room temperature for 15 minutes. Unreacted chemicals were quenched with 1 M ethanolamine (pH 8.5) for 5 minutes. The activated sensors were rinsed with PBS and loaded with 4 µL of 25 µM Bn or anti-GFP Nb, respectively. The coated sensors were washed with PBS, and 2.5 µM GFP-Bs in PBS was introduced to determine binding kinetics. Dissociation was measured by placing the sensor in PBS.

*Preparation of protein conjugated NHS-microspheres*

NHS-activated agarose microspheres were activated according to the manufacturer's instructions. For each type of coating (e.g., control, Nb, Bn, Bs), 100 µL of microspheres (approximately 50% w/w) were coated overnight at 4 °C with 0.5 mL of 25 µM Bn or 25 µM anti-GFP Nb, respectively. Control particles were loaded with 0.5 mL of PBS. All particles were blocked with 0.5 mL of 3% BSA solution, and unreacted NHS groups were quenched with 100 µL of 1 M ethanolamine (pH 8.5) for 5 minutes. The protein-conjugated microspheres were washed twice with PBS and resuspended to a 50% w/w concentration in PBS containing 3% BSA.

*GFP-Bs capture on Bn or antiGFP-Nb coated microspheres*

Bn and anti-GFP-Nb-conjugated microspheres in PBS containing 3% BSA were aliquoted at 50 µL for GFP-Bs capture. The GFP-Bs binding assay was performed by adding 50 µL of 250 nM, 1250 nM, or 2500 nM GFP-Bs sample to the resuspended beads, resulting in final concentrations of 125 nM, 625 nM, and 1250 nM, respectively, in a total volume of 100 µL. After a 5-minute incubation in the dark, the microspheres were centrifuged and washed twice with PBS. The washed microspheres were resuspended in 100 µL of PBS, and fluorescence was measured using a Tecan plate reader at an emission wavelength of 520 nm.

*Secondary nanobody fluorescence labelling*

The NHS-ATTO488 dye was dissolved in DMSO to a concentration of 2 mg/mL. For the labeling reaction, 50 µL of 1 mg/mL nanobody 13A5 was incubated with 3 µL of NHS-ATTO488 and 7 µL of $NaHCO_3$ for 1 hour at room temperature. The labeled product was purified by desalting using a MiniTrap column (GE Healthcare). The labeled nanobody was stored at 4 °C for up to 3 months.

*Detection of Bn-Bs enriched CEA with complementary nanobodies*

The standard detection reaction was set up in vials by mixing 20 µL of 0.1 mg/mL 2D5-Bn, 20 µL of CEA (diluted in PBS to various concentrations), and 20 µL of 0.1 mg/mL ATTO488-labeled 13A5. After incubation in the dark for 10 minutes, the nanobody-CEA complex was captured using 10 µL of 50% w/w Bs-conjugated beads. The microspheres were isolated by centrifugation at 14,800 rpm for 5 minutes and washed twice with PBS. The washed microspheres were resuspended in 100 µL of PBS and transferred to a 96-well microplate for fluorescence measurement.

Alternatively, the nanobody-CEA complex was captured using 10 µL of 50% w/w Bn-conjugated beads from a mixture of 20 µL of 0.1 mg/mL 2D5-Bs, 20 µL of CEA (diluted in PBS to various concentrations), and 20 µL of 0.1 mg/mL ATTO488-labeled 13A5.

The specificity of the designed system was assessed and compared using an excess amount of primary and secondary nanobodies (20 µL of 0.5 mg/mL primary Nb and 20 µL of 0.5 mg/mL 13A5-ATTO488) in PBST containing 3% BSA to detect 20 µL of 1 µg/mL CEA. Non-specific binding between protein components was tested using NHS-blocked, Bn-coated, and Bs-coated beads.

*Fluorescence microscopy*

SUPERFROST® PLUS glass slides (Thermo Scientific) and 0.17 mm coverslips (ZEISS) were used for fluorescence microscopy. Imaging was performed using an epifluorescence microscope (BX51, Olympus) equipped with a 100× NA 1.30 imaging objective (UPLFLN 100X, Olympus). Washed microspheres in a 2 µL volume were imaged with consistent settings: gain (0 dB) and exposure time (500 ms).

*Surface supported sandwich binding of primary Nb, antigen and secondary Nb*

The sandwich binding performance of the 2D5-CEA-13A5 complex was evaluated using a regenerated quartz light interferometer, following a previously published method[2]. Briefly, the quartz surface was regenerated by incubating the biosensor tip in a modified piranha solution (a mixture of $H_2O_2$, water, and ammonia in a 1:3:1 ratio) at 70 °C in a water bath for 30 minutes, followed by rinsing with Milli-Q water. A silica-binding peptide-incorporated 2D5, namely 2D5-CotB1p, was introduced at 5 μM to the regenerated surface, and association occurred spontaneously due to the CotB1p peptide. The 2D5-CotB1p-coated surface was used for kinetic measurements of 1–5 μM CEA, followed by complementary binding with 5 μM of the secondary nanobody 13A5. Data analysis was performed using BLItz Pro 1.3 software.

### Supplementary results

**Supplementary Table S1** List of plasmids used in this work.

| Plasmid name | Plasmid backbone | Encoded protein | Purpose |
|---|---|---|---|
| pMT416 | | Bn | Bn expression |
| pNbBnBs | pETDuet-1 | 2D5-Bn | 2D5-Bn expression |
| pBn2 | pBAD33 | 2D5-Bn | 2D5-Bn for dual transformation |
| pMT316 | | Bs | Bs expression |
| pBs1 | pET28a | Bs | Bs for dual transformation |
| pNbBs | pET28a | 2D5-Bs | 2D5-Bs expression |
| pGFPBs | pET28a | GFP-Bs | GFP-Bs expression |
| pNb1 | pET28a | 2D5 | Control 2D5 expression |
| pNb2 | pET28a | 13A5 | Control 13A5 expression |

**Supplementary Table S2** List of synthesized genes for this work.

| Bn for co-transformation |
|---|
| HHHHHHAQVINTFDGVADYLQTYHKLPDNYITKSEAQALGWVASKGNLADVAPGKSIGGDIFSNRE<br>GKLPGKSGRTWREADINYTSGFRNSDRILYSSDWLIYKTTDHYQTFTKIR |
| Bs for co-expression |
| KKAVINGEQIRSISDLHQTLKKELALPEYYGENLDALWDALTGWVEYPLVLEWRQFEQSKQLTENG<br>AESVLQVFREAKAEGADITIILS |
| GFP-Bs |
| HHHHHHSSGVSKGEELFTGVVPILVELDGDVNGHKFSVSGEGEGDATYGKLTLKFICTTGKLPVPW<br>PTLVTTLTYGVQCFSRYPDHMKQHDFFKSAMPEGYVQERTIFFKDDGNYKTRAEVKFEGDTLVNRI<br>ELKGIDFKEDGNILGHKLEYNYNSHNVYIMADKQKNGIKVNFKIRHNIEDGSVQLADHYQQNTPIG<br>DGPVLLPDNHYLSTQSALSKDPNEKRDHMVLLEFVTAAGITLGMDELYKGGGGSGGGGSKKAVING<br>EQIRSISDLHQTLKKELALPEYYGENLDALWDALTGWVEYPLVLEWRQFEQSKQLTENGAESVLQV<br>FREAKAEGADITIILS |
| 2D5-Bs |
| QVQLQESGGGLVQAGGSLRLSCVASGRTFSSHPMGWFRQAPGKEREFVAGISWSGGSTHYADSVKG<br>RFTISRDTAKNTVYLQMNSLKPEDTAVYYCNAALSERTPIATMPSQYDYWGQGTQVTVSSGGGGSG<br>GGGSKKAVINGEQIRSISDLHQTLKKELALPEYYGENLDALWDALTGWVEYPLVLEWRQFEQSKQL<br>TENGAESVLQVFREAKAEGADITIILSHHHHHH |
| 2D5-Bn |
| MKQSTIALALLPLLFTPVTKAQVQLQESGGGLVQAGGSLRLSCVASGRTFSSHPMGWFRQAPGKER<br>EFVAGISWSGGSTHYADSVKGRFTISRDTAKNTVYLQMNSLKPEDTAVYYCNAALSERTPIATMPS<br>QYDYWGQGTQVTVSSGGGGSGGGGSAQVINTFDGVADYLQTYHKLPDNYITKSEAQALGWVASKGN<br>LADVAPGKSIGGDIFSNREGKLPGKSGRTWREADINYTSGFRNSDRILYSSDWLIYKTTDHYQTFT<br>KIRHHHHHH |

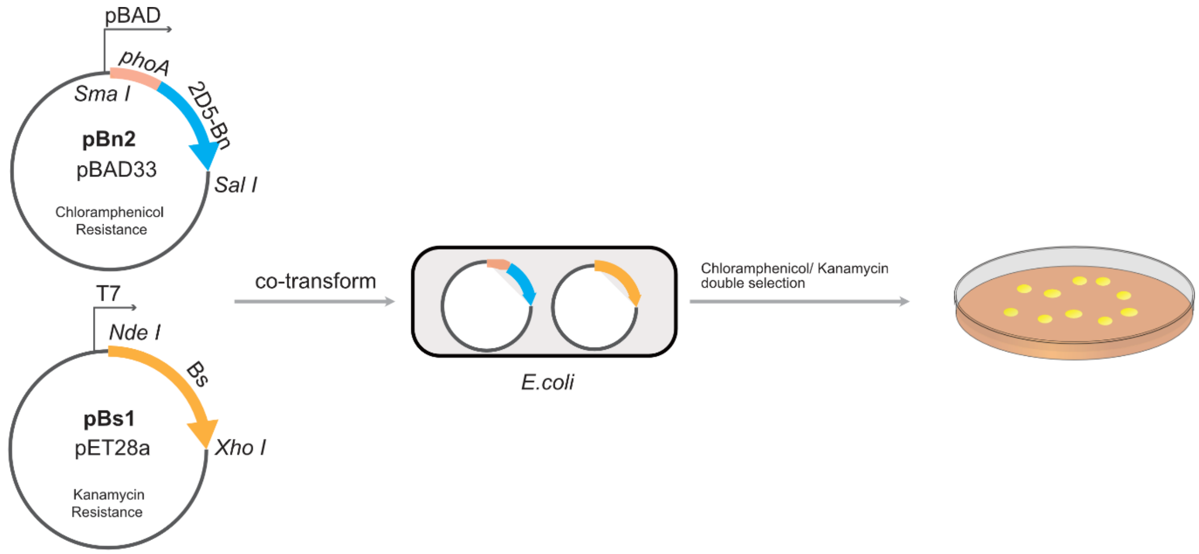


**Supplementary Figure S1. Dual transformation strategy for the expression of the toxic 2D5-Bn gene.** Co-transformation was performed by heat shock at 42 °C for 45 seconds using 50–100 ng of each plasmid. The transformation efficiency was low, with only 1–2 colonies recovered under double selection with kanamycin and chloramphenicol. Although the transformants exhibited protein expression, this strategy was outperformed by the pETDuet-1 plasmid construct due to its complexity.

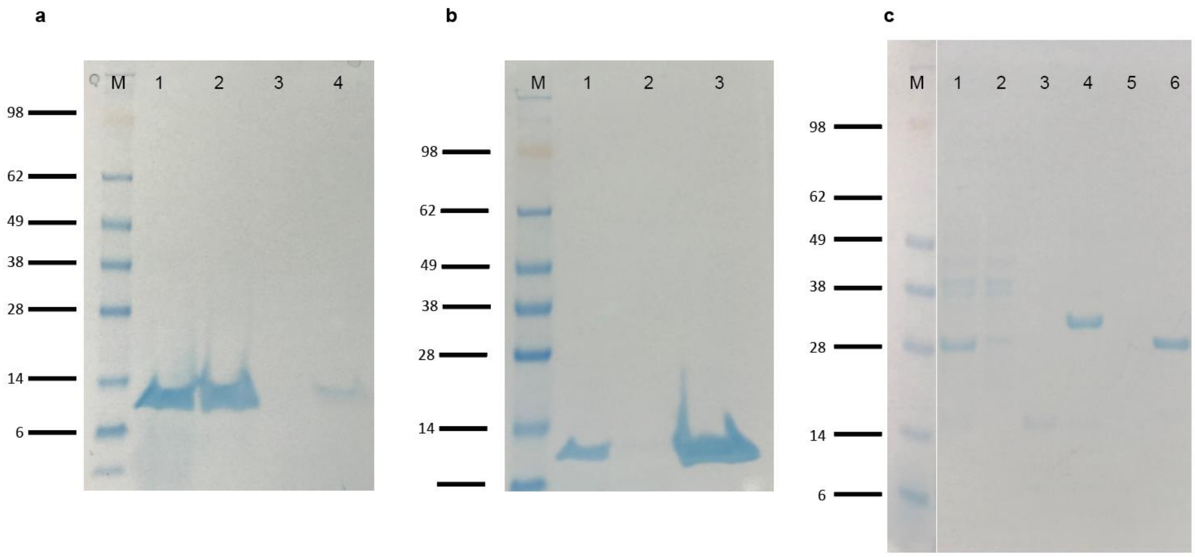


**Supplementary Figure S2. Purification of Bn and 2D5-Bn constructs. a,** Crude Bn extracted with sucrose at 16 mS/cm conductivity did not bind to the HiTrap SP HP cation exchange column. Lane 1: sucrose extract loading; Lane 2: flowthrough; Lane 3: wash; Lane 4: elution. **b,** Dilution of the sucrose extract of Bn to 5 mS/cm conductivity enabled Bn to bind and be purified using the HiTrap SP HP cation exchange column. Lane 1: diluted sucrose extract loading; Lane 2: flowthrough; Lane 3: elution. **c,** Affinity chromatography purification of 2D5-Bn. Lane 1: sucrose extract loading; Lane 2: flowthrough; Lane 3: wash; Lane 4: elution fraction #1; Lane 5: elution fraction #2; Lane 6: elution fraction #3. M: marker.

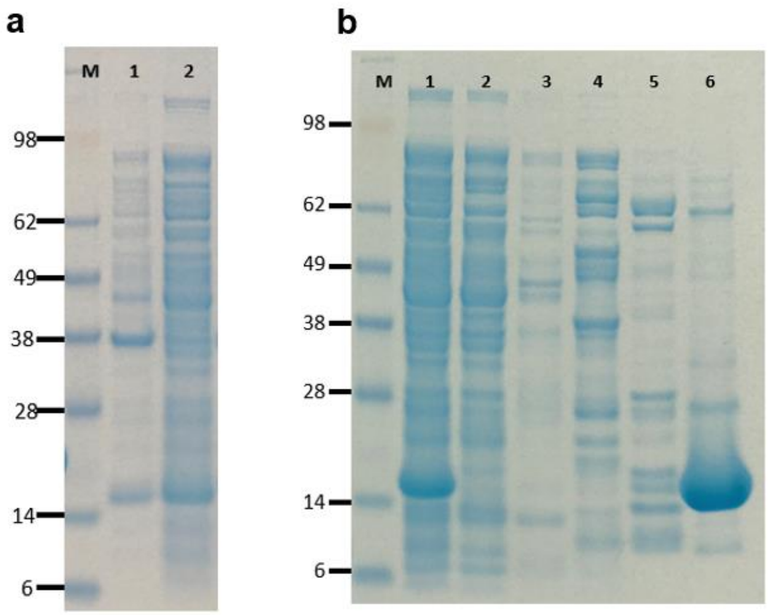


**Supplementary Figure S3. Expression and purification of anti-GFP Nb LaG-16. a,** Expression of anti-GFP Nb LaG-16. M: protein ladder. Lane 1: insoluble fraction of LaG-16 expressed in SHuffle T7 *E. coli* using TB media. Lane 2: soluble fraction of LaG-16 expressed in SHuffle T7 *E. coli* using TB media. **b,** Ni-NTA IMAC purification of LaG-16. Lane 1: soluble fraction loaded. Lane 2: column flowthrough. Lane 3: column wash. Lane 4: pooled elution fractions #1. Lane 5: pooled elution fractions #2. Lane 6: pooled elution fractions #3. The purified protein was desalted into PBS and stored at −80 °C for long-term use.

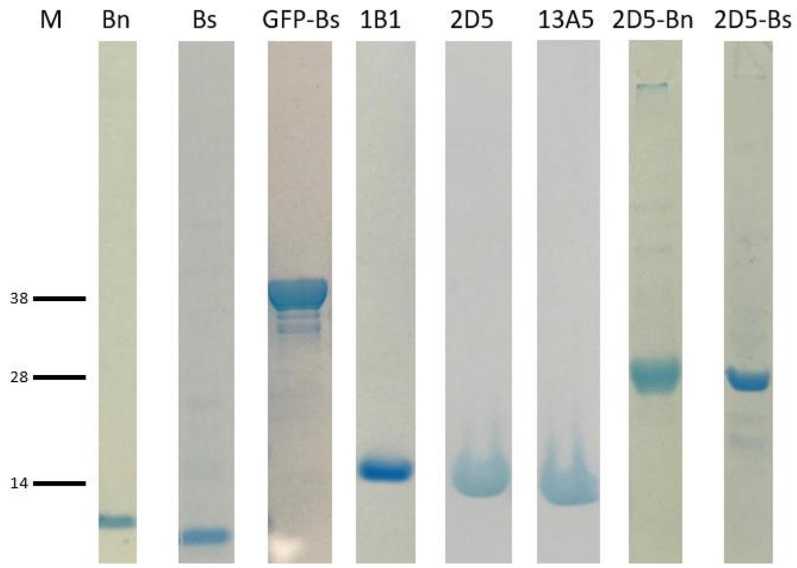


**Supplementary Figure S4. Protein purity of the Bn/Bs toolkit used in this study**. Gel truncates were aligned to the indicated marker lane (M). All proteins were stored in PBS, except for 2D5-Bs, which was stored in 20 mM Tris-HCl (pH 8) and 140 mM NaCl for enhanced long-term stability. Note that the anti-CEA nanobody 1B1 is not included in this work.

**Supplementary Table S3** Sequence verification of Bn, GFP-Bs and 2D5-Bs by mass spectrometry. The sequence of 2D5-Bn was not confirmed by mass spectrometry.

| Description | \|Log Prob\| | Best \|Log Prob\| | Best score | Total Intensity | # of spectra | # of unique peptides | # of mod peptides | Coverage % | # AA's in protein |
|---|---|---|---|---|---|---|---|---|---|
| >gi\|Barnase\|Barnase | 370.96 | 14.76 | 946.40 | 9630985714.5 | 659 | 66 | 0 | 89.09 | 110 |
| >gi\|GFP_barstar\|GFP-barstar | 1066.63 | 24.84 | 1698.40 | 8085521349.7 | 1058 | 156 | 5 | 92.05 | 365 |
| >gi\|2D5_barstar\|2D5-barstar | 919.13 | 17.95 | 1192.30 | 1100781134.5 | 465 | 122 | 12 | 89.20 | 250 |

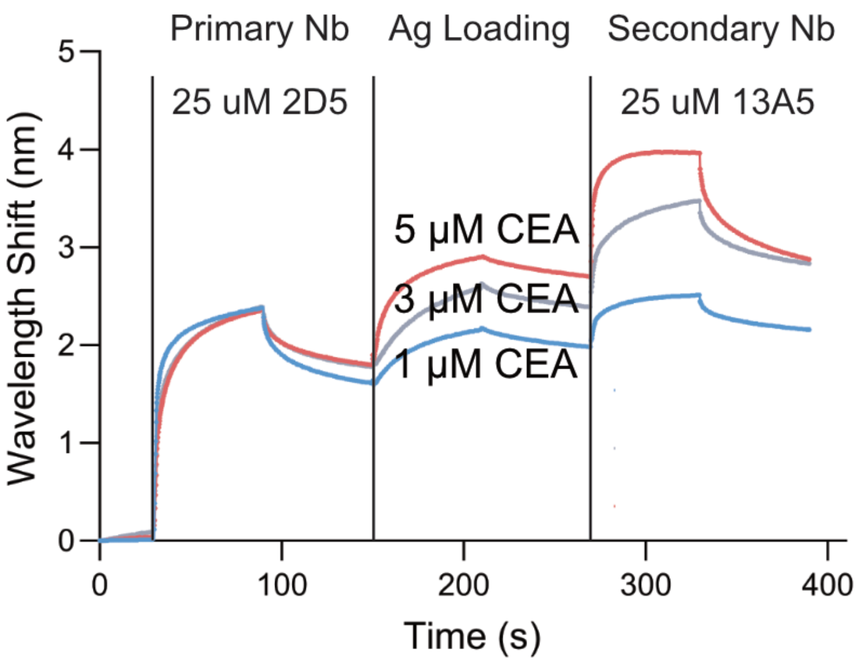


**Supplementary Figure S5. Sandwich binding of primary Nb, antigen, and secondary Nb.** The sandwich binding assay on a regenerated quartz sensor was established in our previous work[2].
